# Cleavage of 14-3-3ε by the enteroviral 3C protease dampens RIG-I mediated antiviral signaling

**DOI:** 10.1101/2022.08.31.505975

**Authors:** Daniel D.T. Andrews, Yasir Mohamud, Marli Vlok, Dorssa Akbari Bani, Brenna N. Hay, Leonard J. Foster, Honglin Luo, Christopher M. Overall, Eric Jan

## Abstract

Viruses have evolved diverse strategies to evade the host innate immune response and promote infection. The RIG-I-like-receptors RIG-I and MDA5 (RLRs) are antiviral factors that sense viral RNA and signal downstream *via* mitochondrial antiviral-signaling protein (MAVS) to activate type I interferon (IFN) expression. 14-3-3ε is a key component of the RIG-I translocon complex that interacts with MAVS at the mitochondrial membrane; however, the exact role of 14-3-3ε in this pathway is not well understood. In this study, we demonstrate that 14-3-3ε is a direct substrate of both the poliovirus and coxsackievirus B3 (CVB3) 3C proteases (3C^pro^), and that it is cleaved at Q236↓G237, resulting in the generation of N- and C-terminal fragments of 27.0 and 2.1 kDa, respectively. Expression of the N-terminal cleavage fragment in cells reduces *IFNB* mRNA production during poly(I:C) stimulation, thus suggesting an antagonistic effect in the presence of the endogenous 14-3-3ε protein. The N-terminal 14-3-3ε fragment does not interact with RIG-I in co-immunoprecipitation assays, nor can it facilitate RIG-I translocation to the mitochondria. Probing the intrinsically disordered C-terminal region identifies key residues responsible for RIG-I signaling. Finally, overexpression of the N-terminal fragment promotes CVB3 infection and influenza A virus (H1N1) RNA production and reduces *IFNB* mRNA production during infection. The strategic enterovirus 3C^pro^-mediated cleavage of 14-3-3ε antagonizes RIG-I signaling by disrupting critical interactions within the RIG-I translocon complex, thus contributing to evasion of the host antiviral response.

**Author Summary:** Host antiviral factors work to sense virus infection through various mechanisms, including a complex signaling pathway known as the RIG-I like receptor (RLR) pathway. This pathway drives the production of antiviral molecules known as interferons, which are necessary to establish an antiviral state in the cellular environment. Key to this antiviral signaling pathway is the small chaperone protein 14-3-3ε, which facilitates the delivery of a viral sensor protein, RIG-I, to the mitochondria. In this study, we show that the enteroviral 3C protease cleaves 14-3-3ε during infection, rendering it incapable of facilitating this antiviral response. We also find that the cleavage fragment inhibits RIG-I signaling and promotes virus infection. Our findings reveal a novel viral strategy that restricts the antiviral host response and provides insights into the mechanisms underlying 14-3-3ε function in RIG-I antiviral signaling.

## Introduction

In response to viral infections, mammalian cells have evolved several distinct mechanisms to sense and respond to the presence of a virus. Sensing of exogenous genetic material, proteins, and other viral markers through these mechanisms in part triggers an enhanced antiviral state in the cell, characterized by expression of interferons (IFNs) and, subsequently, interferon-stimulated genes (ISGs) that restrict infection [1]. In an ‘arms race’ between virus and host, viruses must counter these mechanisms and as such, have evolved intricate evasion mechanisms, such as the capping of viral genetic material to mimic cellular DNA, sequestering of the replication complex to shield the viral particles from detection and the proteolytic processing of cellular antiviral proteins [2–5].

Mammalian cells sense RNA virus infections through membrane-associated toll-like receptors and cytoplasmic retinoic acid-inducible gene I (RIG-I)-like receptors (RLRs) including RIG-I (which senses short double-stranded RNAs (dsRNA)), melanoma differentiation-associated protein 5 (MDA5, which senses long dsRNAs) and laboratory of genetics and physiology 2 (LGP2, which acts as a regulator for other RLRs) [6–8]. In general, RLRs are activated upon the detection of viral dsRNA and RNAs containing a 5’ triphosphate moiety but no methyl cap, though MDA5 can also detect foreign RNAs lacking 5’ triphosphates [1]. Upon detection of viral RNA, RIG-I and MDA5 undergo conformational changes from a “closed” to “open” conformation [9]. The “closed” conformation prohibits access to the Caspase Activation and Recruitment Domains (CARD), while the “open” conformation exposes them. Access to the CARDs is dependent on several factors, such as ubiquitination of RIG-I with K63-polyubiquitin chains and the association of RIG-I with chaperone proteins such as tripartite motif containing 25 (TRIM25) and 14-3-3ε [10,11]. These interactions enable the CARDs of RIG-I to associate with the CARDs of mitochondrial antiviral-signaling protein (MAVS), which serves as an essential step in the RLR pathway [12]. MAVS activation via the CARDs then triggers a downstream cascade that drives type 1 IFN production [6,13]. Notably, the activation of these pathways is virus-specific, with either RIG-I or MDA5 acting as the primary sensor for a given virus during infection [14].

A subset of 14-3-3 proteins play key roles in the RIG-I and MDA-5 signaling pathways. The 14-3-3 family of proteins consists of seven unique isoforms: β, γ, ε, ζ, η, σ, and τ, that function as regulatory molecules in cellular pathways such as cell death and apoptosis, cell cycle regulation, and the cellular antiviral response [15–21]. 14-3-3 proteins adopt a conserved α-helical fold, in which a binding groove recognizes phosphorylated serine or threonine residues within an RSXpSXP or RXXXpSXP motif [16,22]. 14-3-3 interactions regulate protein function by inducing a conformational change, sequestering or relocalizing the target protein, or acting as a molecular scaffold [23,24]. The primary difference between the various 14-3-3 proteins lies in the variable C-terminal region, the function of which is poorly understood. Liu *et al* [11] identified 14-3-3ε as a key interaction partner of RIG-I during activation, demonstrating that 14-3-3ε binding is necessary for the translocation of RIG-I to the mitochondria, which is an essential step in its interaction with and activation of MAVS. In addition to RIG-I interactions with 14-3-3ε and TRIM25 [10], UFL1-mediated ufmylation is a key step in RIG-I translocation [25]. In a comparable manner, highlighting the specificity of 14-3-3 proteins, 14-3-3η promotes MDA5 activation and translocation in HCV-infected cells [15].

Several viruses have evolved countermeasures to this pathway, highlighting the importance of RLRs and 14-3-3 proteins in the cellular antiviral response. The NS3 protein of dengue and Zika virus contains a phosphomimic domain that binds to and sequesters 14-3-3ε, antagonizing the RIG-I response and promoting viral replication [20,26]. The influenza A virus NS1 protein similarly interacts with 14-3-3ε and disrupts RIG-I translocation to the mitochondria [27]. Epstein-Barr virus, Kaposi sarcoma-associated herpesvirus, and human cytomegalovirus all encode ubiquitin deconjugases that interact with 14-3-3 proteins to drive TRIM25 aggregation and inactivation, promoting infection and inhibiting IFN production [28]. Despite being a key RIG-I regulatory factor that is targeted by several distinct viral families, the exact role of 14-3-3ε in RIG-I translocation has not been fully elucidated.

In this study, we showed that 14-3-3ε is cleaved during poliovirus and coxsackievirus B3 (CVB3) infection, and that it is a direct target of the enteroviral 3C protease (3C^pro^) with cleavage occurring between amino acids Q236↓G237 of 14-3-3ε. We demonstrated that expression of the N-terminal cleavage fragment of 14-3-3ε (“3CN”) blocks RIG-I-mediated *IFNB* transcription. Further, we showed that 14-3-3ε 3CN cannot interact with RIG-I, and thereby disrupts the translocation of the RIG-I complex to the mitochondria. Finally, we revealed that the expression of 14-3-3ε 3CN promotes productive infection and restricts *IFNB* production during virus infection. These results reveal a novel strategy by which enteroviruses restrict and evade the host innate immune response.

## Results

### Cleavage of 14-3-3ε in poliovirus and CVB3 infected cells

We previously used Terminal Amine Isotopic Labeling of Substrates (TAILS) to identify host proteins that are cleaved by the enteroviral 3C^pro^ [3,4,29,30]. Mining this TAILS dataset [3] to investigate substrates of poliovirus 3C^pro^, we identified the high-confidence neo-N-termini peptide ^237^GDGEEQNKEA from human 14-3-3ε **(Fig 1A)**, a known chaperone protein and key regulator of the RIG-I activation pathway [11]. Consistent with the preferred Q↓G (P1↓P1’) specificity of 3C^pro^ [31], the TAILS-identified cleavage site in 14-3-3ε is between Q236↓G237, prompting us to further investigate the functional role of cleavage of this protein.

**Figure 1.**
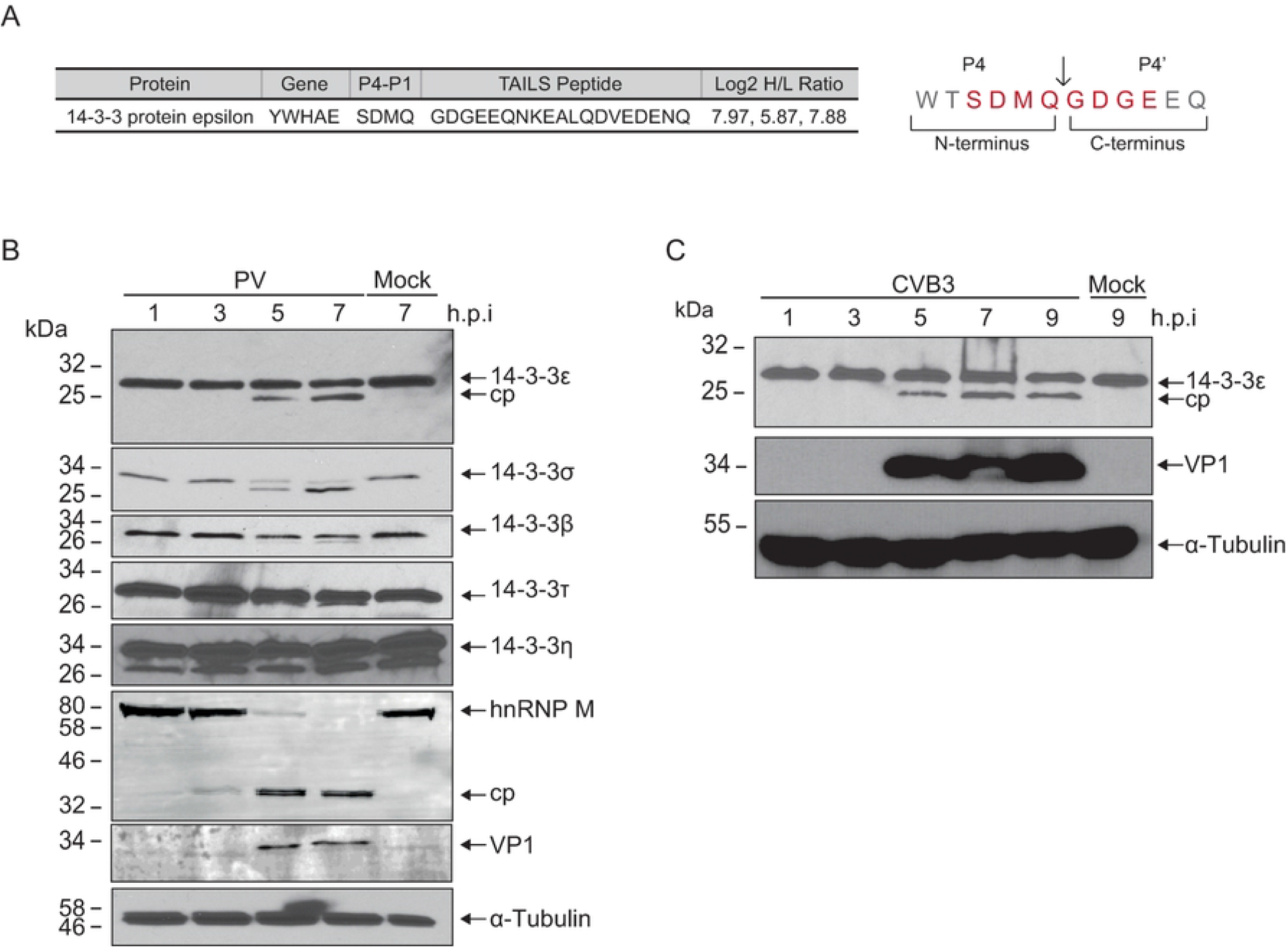
Cleavage of 14-3-3 family members in poliovirus and coxsackievirus B3 infection. (A) High-confidence 14-3-3ε cleavage peptide identified via TAILS analysis. The TAILS peptide was identified *via* mass spectrometry analysis with the upstream P4-P1 and downstream P1’-P4’sites indicated. Log_2_ H/L ratio was determined across three independent experiments (infected cells – heavy, mock – light). (B) Representative immunoblot of poliovirus-infected cell lysates (MOI = 10) infected for the indicated time point. 20 µg total lysate was separated on a 12% SDS-PAGE gel. (C) Representative immunoblot of CVB3-infected cell lysates (MOI = 10) infected for the indicated time point. 20 µg total lysate was separated on a 12% SDS-PAGE gel. “cp” = cleavage product.

First, we sought to determine whether 14-3-3ε is cleaved during virus infection. We infected HeLa cells with poliovirus and probed for 14-3-3ε by immunoblotting. A lower molecular weight product of 14-3-3ε cleavage product was observed 5-hours post-infection (h.p.i.) **(Fig 1B)**. The predicted cleavage event between Q236↓G237 would result in two protein fragments, an N- and C-terminal fragment of ∼27.0 and ∼2.1 kDa respectively. To determine whether cleavage is specific to poliovirus or if it is a general strategy employed during enterovirus infection, we monitored the state of 14-3-3ε in CVB3-infected HeLa cells. Like that observed in poliovirus-infected cells, a cleavage product of 14-3-3ε was observed in CVB3-infected cells **(Fig 1C)**, suggesting a common viral cleavage strategy employed by enteroviruses.

We next examined whether other 14-3-3 family members are cleaved during infection. 14-3-3σ, β, and τ were also cleaved to varying extents during poliovirus infection **(Fig 1B)**. Similar to 14-3-3ε, cleavage fragments of 14-3-3σ and τ were detected as early as 5 h.p.i, while 14-3-3σ was cleaved to completion by 7 h.p.i. By contrast, no detectable 14-3-3η Cleavage products were detected during infection. These results indicated that 14-3-3 proteins are differentially targeted in poliovirus-infected cells.

### Endogenous 14-3-3ε is cleaved directly by poliovirus 3C^pro^

14-3-3ε is a known substrate of caspase-3 during apoptosis and apoptosis can be induced at late stages of poliovirus infection [21,32,33]. To determine whether 14-3-3ε is targeted by caspases in poliovirus-infected cells, we pre-treated cells with zVAD-FMK, a pan-caspase inhibitor, and subsequently infected the cells with poliovirus for 7 hours. As expected, poly (ADP-ribose) polymerase (PARP), a known substrate of caspase-3, was cleaved in poliovirus-infected cells, but was not cleaved in the presence of zVAD-FMK **(Fig 2A)**. By contrast, 14-3-3ε cleavage was still detected in infected, zVAD-FMK-treated cells, thus indicating that 14-3-3ε is not cleaved by caspases in poliovirus-infected cells.

**Figure 2.**
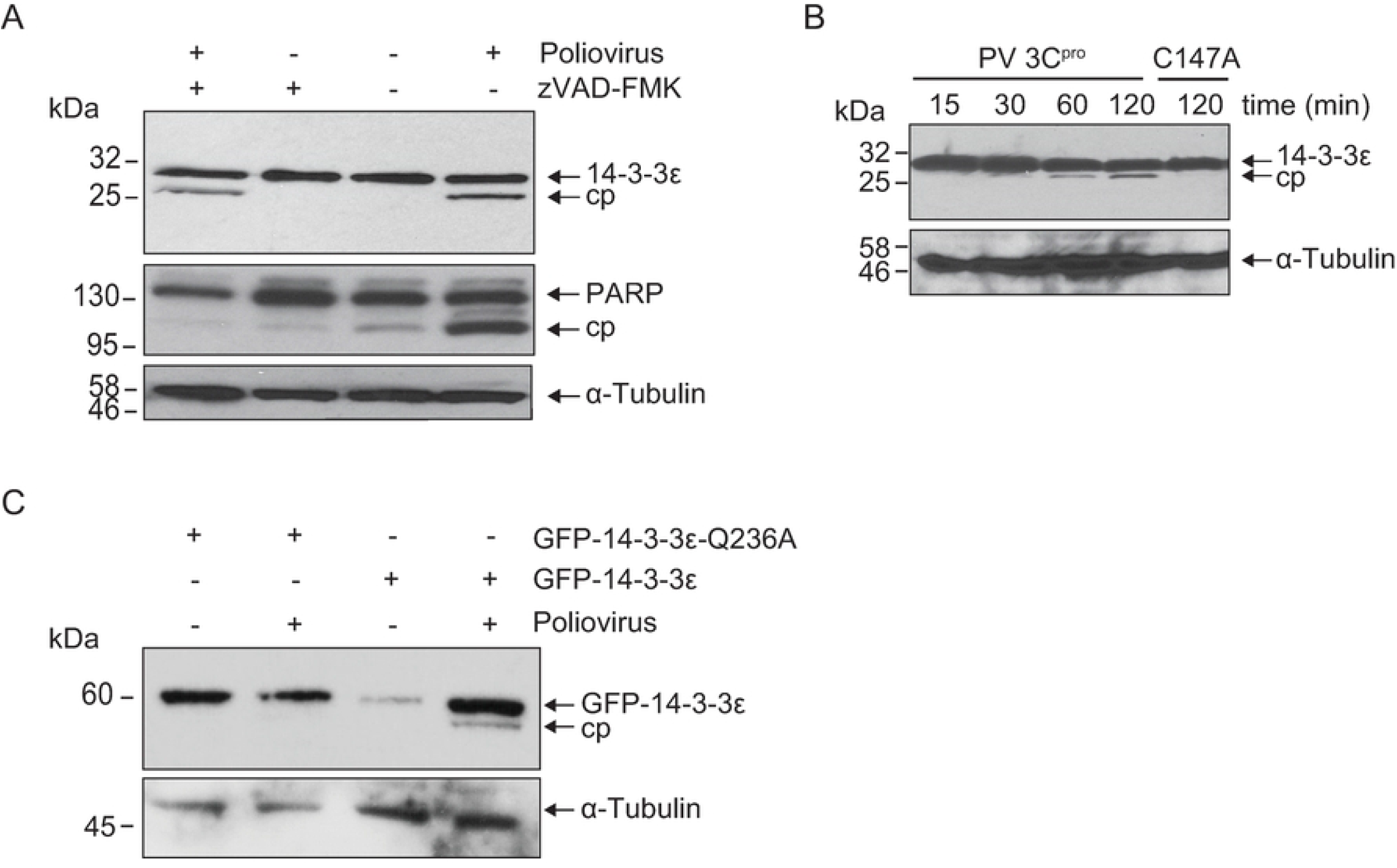
14-3-3ε is cleaved by poliovirus 3C^pro^. (A) Representative immunoblot of poliovirus-infected HeLa cells in the presence or absence of zVAD-FMK at 7 hours post infection (MOI = 10). (B) Representative immunoblot of an *in vitro* cleavage assay performed in HeLa cell lysates. 1 µg clarified cell lysate was incubated with either wild type (WT) or catalytically inactive (C147A) purified recombinant 3C^pro^ (100 ng/µL final concentration). (C) Representative immunoblot of poliovirus-infected HeLa cell lysates transfected with either the 14-3-3ε GFP-WT or the cleavage-resistant mutant, GFP-14-3-3ε (Q236A) (MOI = 10). “cp” = cleavage product.

To determine whether 14-3-3ε is a direct substrate of 3C^pro^, we performed an *in vitro* cleavage assay using HeLa cell lysates incubated with either purified, recombinant wild type (WT) or catalytically inactive (C174A) poliovirus 3C^pro^. As expected, a cleavage fragment of 14-3-3ε, ∼27 kDa, was detected in reactions containing purified wild type, but not the mutant C147A 3C^pro^ **(Fig 2B)**. The ∼27.0 kDa mass of the cleavage fragment was similar to that observed in poliovirus-infected cells (**Fig 1B, 2B**).

The N-terminomics TAILS analysis identified cleavage of 14-3-3ε between Q236↓G237. To confirm this, we generated GFP-tagged 14-3-3ε expression constructs expressing either the wild type 14-3-3ε, or a non-cleavable construct containing a glutamine to alanine mutation at the P1 site (GFP-14-3-3ε(Q236A)). We transfected the wild type or GFP-14-3-3ε(Q236A) constructs into HeLa cells and then infected the cells with poliovirus. Supporting our earlier observations, a cleavage product was detected in poliovirus-infected cells expressing the wild type construct, but not expressing the mutant GFP-14-3-3ε(Q236A) **(Fig 2C)**. In summary, these results demonstrated that 14-3-3ε is a direct substrate of poliovirus 3C^pro^ and that cleavage occurs at the Q236↓G237 site.

### Effects of the 14-3-3ε N-terminal cleavage product on cell viability

A previous report showed that 14-3-3ε is targeted by caspase-3, which then leads to the release of BCL2-associated agonist of cell death to promote apoptosis [21]. Interestingly, the caspase-3-mediated cleavage site occurs at D238, just two amino acids downstream of the Q236 cleavage site of 3C^pro^, and the caspase-induced N-terminal fragment plays a pro-apoptotic role [21]. To determine whether the expression of these N-terminal cleavage fragments of 14-3-3ε have effects on cell viability, we generated a set of expression constructs containing a myc epitope tag at the N-terminal. Alongside the wild type (“WT”), we generated constructs that terminated at either Q236 (“3CN”) or D238 (“CaspN”) to mimic that of a stable cleavage fragment **(Fig 3A)**. Following transfection, we first assessed, by immunoblotting, caspase activity *via* PARP cleavage **(Fig S1A)** [34]. In all cases, the expression of the tagged 14-3-3ε proteins did not lead to detectable cleavage or loss of full-length PARP **(Fig. S1A)**. Furthermore, overexpression of these 14-3-3ε proteins did not affect cell viability **(Fig. S1B)**. Collectively, these results demonstrated that expression of the truncated 14-3-3ε protein fragments does not activate caspases nor induce cell death in A549 cells.

**Figure 3.**
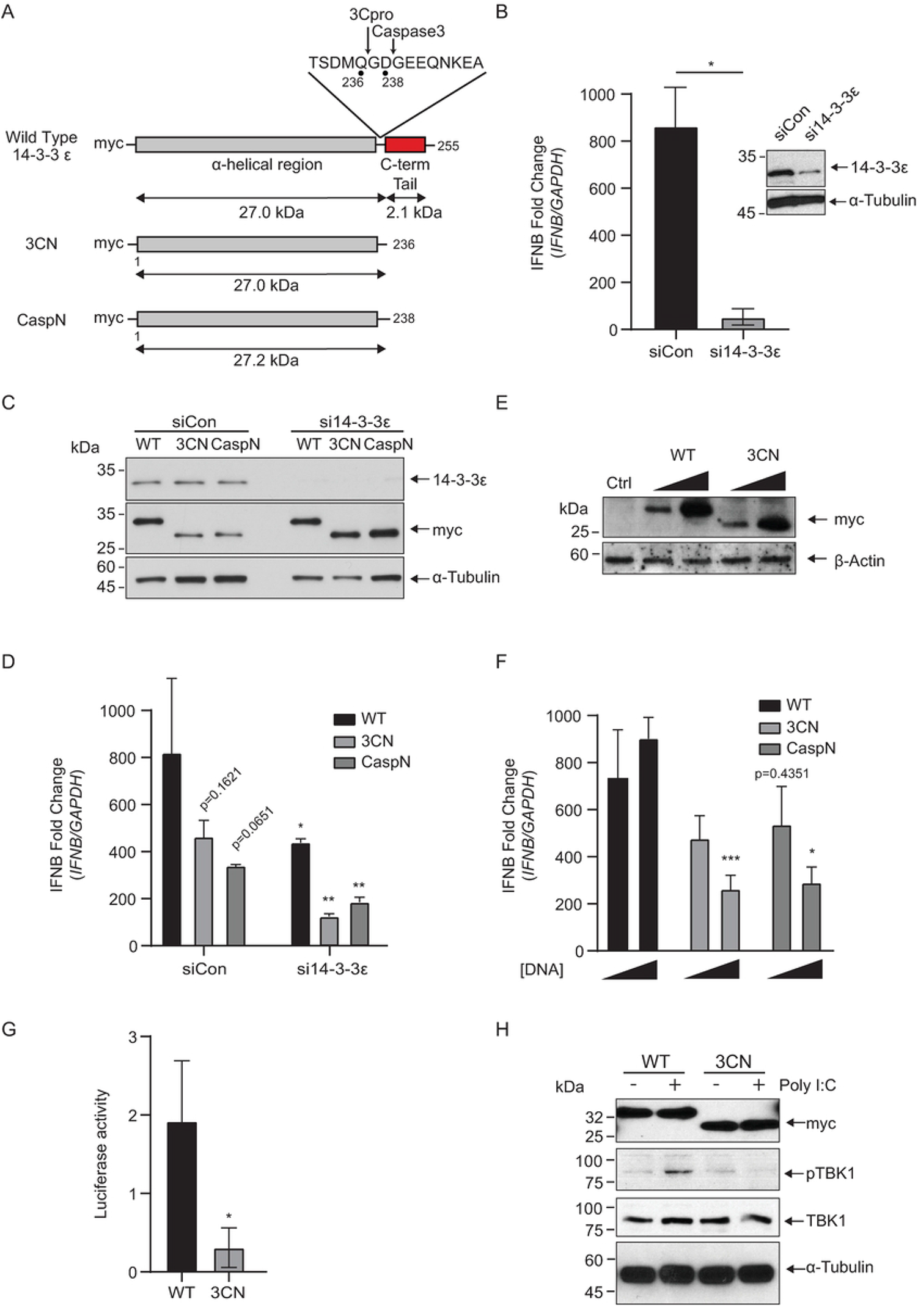
N-terminal 14-3-3ε dampens RLR-stimulated *IFNB* mRNA production. (A) Schematic of 14-3-3ε expression constructs. Wild type (“WT”) contains the N-terminal α-helical region (grey) and the C-terminal tail (red). Inset: amino acids flanking the 3C^pro^ cleavage site (Q236) and the caspase-3 cleavage site (D238). “3CN” = 3C^pro^ N-terminal cleavage product (middle); “CaspN” = caspase-3 N-terminal cleavage product (bottom). (B) RT-qPCR of *IFNB* mRNA from cells treated with either siCon or si14-3-3ε for 24 hours and poly(I:C) for 6 hours. Expression levels were normalized internally to *GAPDH* and fold changes are relative to unstimulated control samples. Inset: immunoblot representation of lysates treated with siCon or si14-3-3ε for 24 hours. * = p<0.05 (p=0.0179). (C) Immunoblot analysis of A549 cells treated with either siCon or si14-3-3ε for 24 hours and transiently transfected with the indicated DNA for a further 24 hours. (D) RT-qPCR of *IFNB* mRNA from cells treated with either siCon or si14-3-3ε for 24 hours, the indicated DNA for a further 24 hours, and poly(I:C) for 6 hours. Expression levels were normalized internally to *GAPDH* and fold changes are relative to unstimulated control samples. * = p<0.05; ** = p<0.005 compared to siCon equivalent (WT, p=0.0418; 3CN, p=0.0011; CaspN, p=0.0014). (E) Immunoblot analysis of A549 cells transfected with an increasing dose of either WT or 3CN 14-3-3ε overexpression constructs (100 ng or 500 ng) for 24 hours. (F) RT-qPCR of *IFNB* mRNA from cells transiently transfected with the indicated DNA for 24 hours and poly(I:C) for 6 hours. Expression levels were normalized internally to *GAPDH* and fold changes are relative to unstimulated control samples. * = p<0.05, *** = p<0.0005 compared to the wild type control (3CN, p=0.0002; CaspN, p=0.0123). (G) Luciferase assay of cells transfected with the indicated DNA for 24 hours and transfected with poly(I:C) for 24 hours. Luminescence detected is the Lucia luciferase protein under the control of an ISG54 promoter and 5xISRE. * = p<0.05 relative to WT control (3CN, p=0.00802). (H) Immunoblot analysis of A549 cells transfected with either WT or 3CN 14-3-3ε constructs for 24 hours and transfected with poly(I:C) for 6 hours. All statistical analyses were performed in GraphPad Prism 9.0; n=3.

### N-terminal 14-3-3ε dampens *IFNB* mRNA production

14-3-3ε is a key component of the RIG-I translocon complex, serving as a chaperone to promote RIG-I signaling and activate the downstream interferon response [25]. We hypothesized that cleavage of 14-3-3ε by 3C^pro^ and the resulting N-terminal fragment disrupts RIG-I signaling. We first determined whether endogenous 14-3-3ε is necessary for *IFNB* mRNA production in A549 cells by transfecting an siRNA targeting the 3’ UTR of the *YWHAE* transcript. Transfection of the 14-3-3ε siRNA in A549 cells resulted in significant knockdown of the endogenous protein by immunoblotting **(Fig 3B**, inset). To activate the RLR pathway, we transfected A549 cells with the dsRNA analogue polyinosinic:polycytidylic acid (poly(I:C)) and then measured *IFNB* mRNA levels by RT-qPCR. Transfection of poly(I:C) in the presence of a control siRNA targeting firefly luciferase (siCon) resulted in an ∼800-fold increase in *IFNB* mRNA levels compared to untreated cells, confirming the activation of the RLR pathway. By contrast, poly(I:C) treatment with an siRNA targeting *YWHAE* significantly reduced the amount of *IFNB* mRNA produced, demonstrating the requirement of 14-3-3ε in RLR pathway activation as previously reported **(Fig 3B)** [11].

Utilizing an siRNA targeting the 3’ UTR of the *YWHAE* transcript allows for exogenous expression of N-terminal 14-3-3ε fragments and their effects on the RLR signaling pathway. Specifically, myc-tagged wild-type, 3CN, and CaspN 14-3-3ε were stably expressed in both control cells and cells depleted of endogenous 14-3-3ε by siRNA treatment **(Fig 3C)**. While the knockdown of 14-3-3ε reduced the production of *IFNB* mRNA **(Fig 3B)**, we observed a moderate rescue of *IFNB* levels in cells transfected with the wild-type 14-3-3ε construct **(Fig 3D)**. By contrast, in si14-3-3ε-treated cells, expression of both 3CN and CaspN 14-3-3ε failed to rescue *IFNB* mRNA production, indicating that the N-terminal 14-3-3ε cleavage fragments cannot support RLR signaling **(Fig 3D)**.

Interestingly, in cells treated with control siRNA, expression of the 3CN and CaspN 14-3-3ε constructs resulted in a markedly lower *IFNB* fold change relative to that of wild-type 14-3-3ε-transfected cells, suggesting that the overexpression of the N-terminal 14-3-3ε cleavage fragments antagonizes RLR signaling, even in the presence of endogenous 14-3-3ε **(Fig 3D)**. To investigate this further, we transfected increasing amounts of each expression construct in A549 cells followed by poly(I:C) stimulation **(Fig. 3E, 3F)**. Overexpression of both 3CN and CaspN 14-3-3ε resulted in a significant decrease in *IFNB* production in a dose-dependent manner **(Fig 3F)**. To further confirm this observation, we monitored RLR signaling using an A549 cell line engineered to express an interferon stimulated response element (5xISRE)-luciferase reporter. Overexpression of 3CN 14-3-3ε reduced relative luciferase activity compared to the wild type 14-3-3ε, providing further confirmation that expression of the N-terminal 14-3-3ε cleavage products can antagonize RLR signaling in A549 cells **(Fig 3G)**.

The 14-3-3ε-containing translocon complex signals upstream of MAVS, which subsequently activates TANK binding kinase (TBK1) phosphorylation [35]. To determine the effects of our 3CN 14-3-3ε expression in RLR signaling, we monitored TBK1 phosphorylation (p-TBK1) by immunoblotting. As expected, poly(I:C) treatment resulted in an increase in p-TBK1 levels in cells transfected with wild-type 14-3-3ε, indicating activation of the RLR signaling pathway **(Fig 3H)**. By contrast, cells expressing 3CN 14-3-3ε led to a notable loss of TBK1 phosphorylation, indicating that expression of 3CN 14-3-3ε dampens the RLR signaling pathway upstream of TBK1 phosphorylation. In summary, these results demonstrate that the truncated, N-terminal fragment derived from 3C^pro^-mediated cleavage of 14-3-3ε can act in a dominant negative manner to block RLR signaling.

### Key residues in the C-terminal tail of 14-3-3ε for RIG-I signaling

Our results indicated that cleavage of 14-3-3ε by 3C^pro^ results in the generation of a truncated N-terminal cleavage fragment that antagonizes RLR signaling. These results also suggested that the C-terminal residues of 14-3-3ε may have an important role in RLR signaling. The C-terminal tail is highly variable among the 14-3-3 family proteins and is predicted to be flexible and disordered [36]. While highly variable between 14-3-3 protein family members, the C-terminus of 14-3-3ε is 100% conserved among mammals and the 3C^pro^-induced QG cleavage site is 100% conserved **(Fig S2A, S2B)**. To date, a complete crystal structure of 14-3-3ε resolving the C-terminal tail has yet to be determined. It has been proposed that charged residues in the C-terminal tail may regulate 14-3-3 function by interacting with the phospho-binding pocket of 14-3-3 proteins [36–38], though the detailed functions of the C-terminal tail of 14-3-3 proteins are not fully understood. Modeling of the structure of the C-terminal domain of 14-3-3ε using RoseTTA fold [39] *in silico* predicted a short, alpha-helical fold **(Fig. 4A)**.

**Figure 4.**
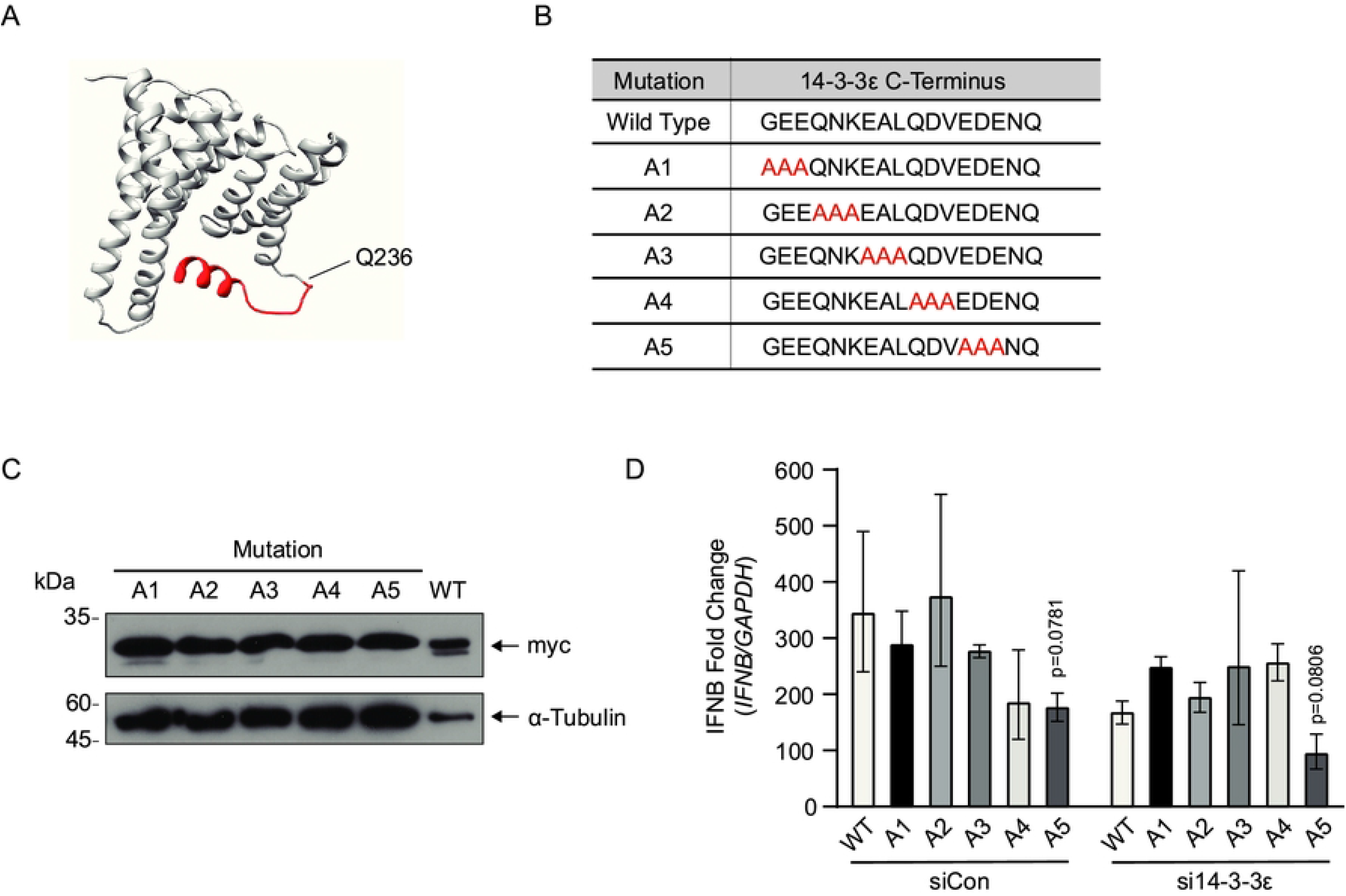
Key residues in the C-terminal region of 14-3-3ε for RIG-I signaling. (A) *In silico* 3D structural model of 14-3-3ε generated using RoseTTA fold and existing crystal structures (PDB: 6EIH). The 14-3-3ε 3C^pro^ cleavage site is indicated (Q236) and the C-terminal fragment is shown in red. (B) Scanning alanine mutagenesis of the 14-3-3ε C-terminus. G237-Q255 were sequentially mutated in codon triplets; mutations indicated in red. (C) Representative immunoblot of the scanning alanine mutants indicated in B. Cells were transfected with the indicated construct for 24 hours. (D) RT-qPCR of *IFNB* mRNA from cells treated with either siCon or si14-3-3ε for 24 hours, the indicated DNA for a further 24 hours, and poly(I:C) for 6 hours. Expression levels were normalized internally to *GAPDH* and fold changes are relative to unstimulated control samples. All statistical analyses were performed in GraphPad Prism 9.0; n=3.

To determine residues within the C-terminal region of 14-3-3ε that may be required for RLR signaling, we used a scanning alanine mutagenesis approach, systematically mutating three consecutive residues to alanine from G237 to Q255 **(Fig 4B)**. We overexpressed alanine-mutated 14-3-3ε in A549 cells **(Fig 4C)** treated with either a control siRNA or si14-3-3ε to address the direct effects of C-terminal 14-3-3ε mutations on RLR signaling. As expected, poly(I:C) treatment increased *IFNB* mRNA levels in siCon-treated cells transfected with the wild-type 14-3-3ε construct **(Fig 4D)**. Similarly, overexpression of 14-3-3ε mutants (A1-A4) resulted in an increase in *IFNB* mRNA levels similar to wild-type 14-3-3ε, suggesting that these residues are not critical for RLR signaling. By contrast, transfection of the 14-3-3ε A5 mutant led to reduced *IFNB* mRNA levels compared to wild-type 14-3-3ε expression in poly(I:C)-stimulated cells. We next determined the effects of these 14-3-3ε mutants in cells depleted of endogenous 14-3-3ε by siRNA treatment. Expression of 14-3-3ε mutants (A1-A4) in the presence of si14-3-3ε exhibited a similar fold change to that of the wild-type 14-3-3ε expressing cells. Transfection of the 14-3-3ε A5 mutant construct in si14-3-3ε-treated cells did not fully rescue *IFNB* mRNA production **(Fig 4D)**. These results suggested that residues 251-253 (E251, D252, E253) of 14-3-3ε are important for RLR signaling.

### N-terminal 14-3-3ε disrupts the RIG-I translocon

Our results showed that overexpression of the 3CN construct disrupted the RLR signaling pathway upstream of TBK1 phosphorylation **(Fig 3H)**. We next determined whether the expression of 3CN 14-3-3ε blocks RIG-I translocation to the MAVS at the mitochondria. To address this, we co-transfected FLAG-RIG-I and either the myc-tagged wild-type or 3CN 14-3-3ε constructs in 293T cells, and then monitored RIG-I localization in the mitochondrial and cytoplasmic fractions by immunoblotting. Confirming membrane fractionation, the mitochondrial marker voltage-dependent anion-selective channel protein 1 (VDAC1) and the cytoplasmic β-actin were enriched in their respective subcellular compartments **(Fig 5A)**. Consistent with previous reports [11], poly(I:C) treatment of cells expressing the wild-type 14-3-3ε construct resulted in an enrichment of FLAG-RIG-I in the mitochondrial fraction. Conversely, expression of 3CN 14-3-3ε caused a loss of enrichment of FLAG-RIG-I in the mitochondrial fraction **(Fig 5A)**. Quantification of the immunoblots confirmed that the amount of FLAG-RIG-I localized to the mitochondria was decreased in cells expressing 3CN 14-3-3ε **(Fig 5B)**. These results indicated that expression of 3CN 14-3-3ε blocks signaling at or upstream of the RIG-I translocation step.

**Figure 5.**
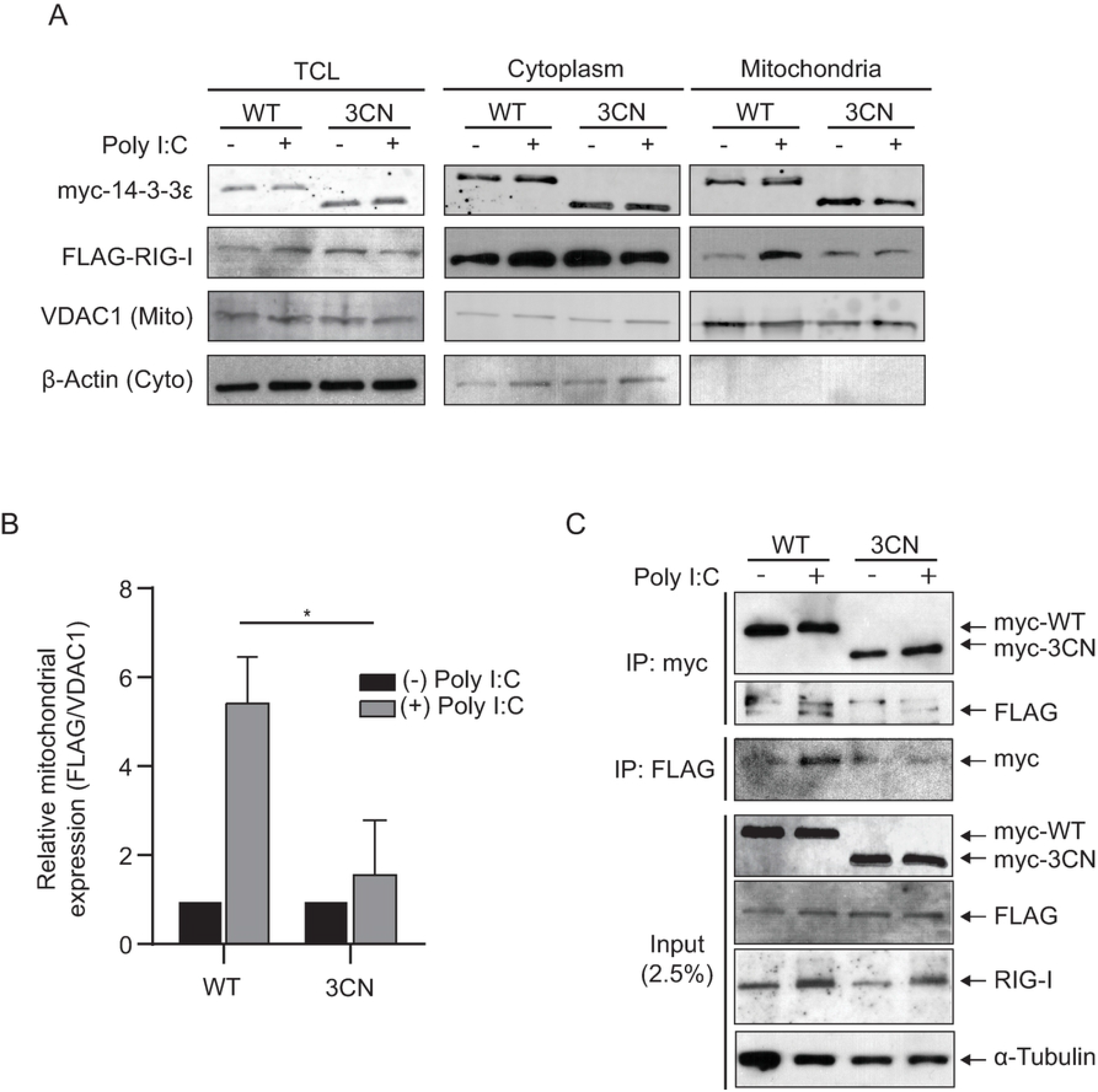
N-terminal 14-3-3ε overexpression disrupts the RIG-I translocon. (A) Immunoblot of myc-14-3-3ε, FLAG-RIG-I, VDAC1, and β-actin before and after subcellular membrane fractionation. Cell lysates were transiently transfected with FLAG-RIG-I and either myc-tagged WT or 3CN 14-3-3ε, constructs for 24 hours, then transfected with poly(I:C) for 6 hours. Lysates were subjected to membrane fractionation and analyzed via Western blot. TCL = total cell lysate; 5% input. (B) Quantification of FLAG and VDAC1 band intensity in mitochondrial fraction of (A). Bands were quantified using ImageJ and FLAG band intensity was normalized to VDAC1 band intensity (n=2). * = p<0.05; p=0.0313. (C) Co-immunoprecipitation assay of cells transiently transfected with FLAG-RIG-I and either myc-tagged wild-type (WT) or 3CN 14-3-3ε constructs for 24 hours, then transfected with poly(I:C) for 6 hours. Lysates were incubated with either myc or FLAG antibody as bait and precipitated using magnetic protein G agarose; 2.5% input. Representative blots are shown from at least three independent experiments.

The RIG-I translocon complex involves several multi-protein interactions that contain at least 14-3-3ε, TRIM25, UFL1 and RIG-I [10,11,25]. To determine whether 3CN 14-3-3ε can interact with RIG-I in poly(I:C)-stimulated cells, we performed co-immunoprecipitation assays in cells expressing FLAG-RIG-I and either the myc-tagged wild-type or 3CN 14-3-3ε proteins **(Fig 5C)**. Expression of tagged RIG-I and the wild-type 14-3-3ε in poly(I:C)-stimulated cells resulted in a detectable interaction between the two proteins (**Fig 5C**). Specifically, poly(I:C) stimulated cells displayed an enrichment of FLAG-RIG-I co-precipitating with the myc-tagged wild-type 14-3-3ε. In the reciprocal immunoprecipitation, the wild-type 14-3-3ε protein co-precipitated with FLAG-RIG-I. Conversely, the 3CN 14-3-3ε protein failed to co-precipitate with FLAG-RIG-I in poly(I:C) stimulated cells **(Fig 5C)**. Together, these results indicated that the N-terminal 14-3-3ε cleavage protein cannot interact with RIG-I in poly(I:C)-stimulated cells.

### N-terminal 14-3-3ε cleavage fragment modulates virus infection

We next addressed whether the N-terminal 14-3-3ε cleavage fragment can modulate virus infection, notably CVB3 and influenza A virus (IAV, H1N1 strain) infections. IAV induces a strong interferon response, while CVB3 targets several factors within the RIG-I signaling pathway [14,40]. First, we investigated the role of 14-3-3ε in CVB3 infection. Depleting endogenous 14-3-3ε resulted in a ∼2 log fold decrease in viral titer, indicating a pro-viral role for 14-3-3ε in CVB3 infection **(Fig 6A)**. We then investigated whether 14-3-3ε contributes to IAV infection. Depletion of 14-3-3ε in A549 cells resulted in a significant decrease in IAV RNA accumulation **(Fig 6B)**. Depleting 14-3-3ε also significantly reduced viral nucleocapsid protein expression **(Fig 6C)**. These results indicated that, like CVB3, 14-3-3ε is required to promote productive IAV infection.

**Figure 6.**
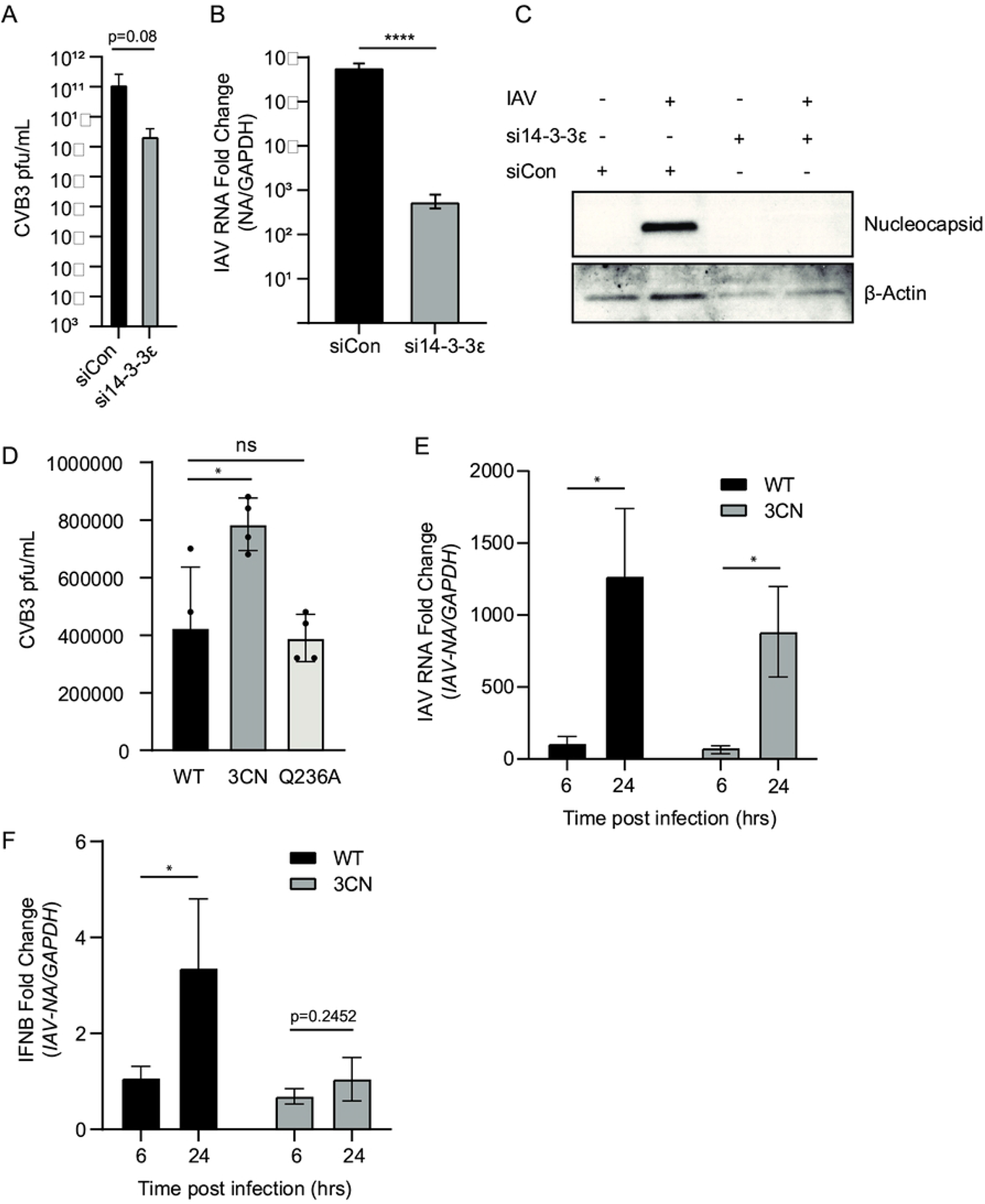
N-terminal 14-3-3ε modulates CVB3 and IAV infection. (A) Viral yield of CVB3-infected A549 cells (MOI=0.01, 24 hours) that were transfected with the indicated siRNAs. Cells were transfected with siRNAs for 48 hours prior to infection. Viral titers were determined (pfu/mL) via plaque assay; n=3, n.s. = not significant (p=0.0753). (B) RT-qPCR analysis of IAV viral RNA from cells transfected with either siCon or si14-3-3ε for 24 hours and subsequently infected with IAV for 24 hours (MOI = 0.01). **** = p<0.00005 compared to siCon (p=1.01×10^−7^). (C) Immunoblot analysis of A549 cells transfected with either siCon or si14-3-3ε for 24 hours and infected with IAV for 24 hours (MOI = 0.01). (D) Viral yield of CVB3-infected HeLa cells (MOI 0.01, 24 hours) transfected with either the wild-type (WT), 3CN, or cleavage-resistant mutant 14-3-3ε (Q236A) overexpression constructs. Cells were infected with CVB3 (MOI 0.01) for 24 hours and viral yield was measured by plaque assay (pfu/mL). n=4. (E, F) RT-qPCR analysis of viral RNA (E) and *IFNB* (F) RNA from A549 cells transfected with either wild-type or 3CN 14-3-3ε constructs for 24 hours, and subsequently infected with IAV (MOI =0.01) for the indicated time points. Expression levels were normalized internally to *GAPDH* and fold changes are relative to uninfected control samples. * = p<0.05; n=7.

To determine whether overexpression of the 3CN 14-3-3ε affected CVB3 infection, and whether preventing cleavage was detrimental, we transfected cells with either the wild-type, 3CN, or the cleavage resistant 14-3-3ε(Q236A) constructs. A549 cells were infected with CVB3 at an MOI of 0.01 for 24 hours, then plaque assays were performed to determine viral yield. There was no difference in CVB3 viral titers in cells transfected with 14-3-3ε(Q236A) compared to that of the wild-type 14-3-3ε construct (**Fig. 6D**). Interestingly, transfection with the 3CN 14-3-3ε construct increased viral yield compared to the wild-type 14-3-3ε transfected cells **(Fig 6D)**, suggesting a pro-viral role of the N-terminal 14-3-3ε cleavage fragment.

We then examined the effects of expressing the wild-type or 3CN 14-3-3ε in IAV-infected cells. We infected cells with IAV at an MOI of 0.1 and measured both *IFNB* and viral RNA levels at 6 and 24 hours post infection by qRT-PCR. Viral RNA levels were similar in cells transfected with either construct **(Fig 6E)**. However, cells transfected with the 3CN 14-3-3ε construct led to reduced *IFNB* mRNA levels compared to that transfected the wild-type 14-3-3ε construct **(Fig 6F)**, consistent with dampening of the RIG-I signaling by the N-terminal 14-3-3ε cleavage protein. These results indicate that expression of the 3CN 14-3-3ε promotes CVB3 infection and reduces both IAV- and poly(I:C)-stimulated RLR signaling.

## Discussion

The RIG-I signaling pathway is a critical first-line host defense in detecting RNA virus infection and activating the antiviral interferon response [41]. Highlighting the importance of this pathway are the diverse viral mechanisms that target RLR signaling to restrict interferon production [18,20,27,28]. 14-3-3ε is a key factor in the RIG-I translocon complex that signals to MAVS, however, the exact role of 14-3-3ε in the translocon complex is not well understood [11]. In this study, we identified 14-3-3ε as a direct target of enterovirus 3C^pro^, which cleaves between Q236 and G237, thus providing insights into the key regions of 14-3-3ε important for RIG-I signaling. Expression of a 3C^pro^-mediated N-terminal 3CN 14-3-3ε cleavage fragment antagonizes RIG-I signaling in a dominant-negative manner and is unable to interact with RIG-I. We further mapped key residues in the C-terminal region of 14-3-3ε that are important for RIG-I signaling, thus providing a mechanistic rationale for cleavage by 3C^pro^ and release of the C-terminal tail. Finally, we showed that the overexpression of the N-terminal fragment promoted CVB3 infection, supporting a pro-viral role for 14-3-3ε cleavage and the N-terminal fragment itself. We propose that strategic cleavage of 14-3-3ε by enterovirus 3C^pro^ contributes to evasion of the host antiviral RIG-I signaling pathway and promoting infection.

How does the 3C^pro^-mediated cleavage of 14-3-3ε disrupt and antagonize RIG-I signaling? 14-3-3ε, along with RIG-I and TRIM25, form a translocon complex that plays a critical role in signalling from RIG-I to MAVS at the mitochondrial membrane [10,11]. Recently, it has been shown that 14-3-3ε is also ufmylated by UFL1 during RIG-I signalling, a further requirement for translocon formation and translocation [25]. Indeed, post-translational modifications of RIG-I, MDA5 and their interaction partners play prominent roles in 14-3-3ε activation and regulation of RLR signalling [16]. The dynamics and interplay of these factors are not fully understood. The N-terminal regions of 14-3-3ε family members contain dimerization sites that mediate homo-or hetero-dimerization with itself or other 14-3-3 proteins, as well as a binding groove that facilitates binding to phosphorylated substrates [16,36]; however, previous studies have shown that 14-3-3ε likely binds to RIG-I in a phosphorylation-independent manner [11], suggesting that 14-3-3ε may interact with RIG-I via an atypical manner or indirectly via other translocon complex proteins. Although the truncated N-terminal fragment does not bind to FLAG-RIG-I **(Fig 5B)**, it is possible that it may also disrupt other interactions within the translocon complex, such as with TRIM25 and UFL1, or some combination of these. Our results also point to some key acidic residues of the C-terminus that potentially promote RIG-I signaling **(Fig 4D)**, thus providing a mechanistic explanation as to why cleavage by 3C^pro^ after Q236 may be strategic to block signaling activity.

As the C-terminus of 14-3-3ε is poorly characterized [36], it remains to be determined how the loss of amino acids 251-253 of 14-3-3ε may abolish functionality. Notably, it has been proposed that the disordered C-terminal region of 14-3-3 proteins may act in an autoinhibitory manner; Truong et al [42] proposed that the C-terminal region was responsible for preventing promiscuity and fine-tuning specific protein-protein interactions. It follows that truncation of the C-terminal tail by the enterovirus 3C^pro^ may alter the interactions between 14-3-3ε and its ligands and disrupt downstream signaling pathways. It is possible that truncation of 14-3-3ε by enterovirus 3C^pro^ does not result in a loss-of-function of 14-3-3ε activity, but rather that the N-terminal fragment is endowed to bind to other cellular proteins. Intriguingly, although our study showed that the N-terminal 14-3-3ε cleavage product cannot interact with RIG-I, expression of the N-terminal 14-3-3ε can promote CVB3 infection, thus pointing to a role in a specific step in the viral life cycle (**Fig 6**). Moreover, depletion studies revealed a critical role of 14-3-3ε In CVB3 infection, thus further indicating a prominent role of the cleavage product(s) of 14-3-3ε in promoting infection. Future studies revealing the interactome of the N-terminal 14-3-3ε cleavage fragment should provide insights into this mechanism.

In enterovirus infections, several proteins in the RLR signaling pathway are cleaved and/or degraded such as RIG-I [40], MDA5, and MAVS [43]. As such, although 14-3-3ε may not be cleaved to completion during infection **(Fig 1B)**, enteroviruses target multiple factors to ensure disabling the RLR signaling pathway. Interestingly, our results strongly suggest that the N-terminal 14-3-3ε cleavage fragment acts in a dominant manner to block RIG-I signaling **(Fig 3, 6)**. However, our results also showed that depletion of endogenous 14-3-3ε is detrimental to CVB3 and IAV infection **(Fig 6)**, which may suggest that the full-length 14-3-3ε protein may retain pro-viral roles. Given that 14-3-3ε normally acts as a chaperone protein for a wide range of signaling pathways and critical interactions with other proteins, it is reasonable to assume that depleting 14-3-3ε would disrupt other cellular functions in such a way that restricts viral replication. Alternatively, as discussed above, the N-terminal 14-3-3ε cleavage product may have other roles in the viral life cycle in promoting infection.

Our results showed that 14-3-3ε is necessary for IAV H1N1 infection (**Fig 6**). This result is in contrast with previous findings that showed that depleting 14-3-3ε promotes IAV infection [27]. We note that our experimental conditions differed from the established literature; Tam *et al* [27] performed their knockdown experiments using a modified virus with a deletion of the NS1 effector domain (ED) and using a higher MOI. Furthermore, Ayllon *et al* [44] speculated that the NS1 ED was responsible for shielding dsRNA from detection by PRRs such as RIG-I; as such, we cannot rule out that the deletion of the NS1 ED may mask critical interactions of RIG-I with 14-3-3ε.

Targeting host cellular proteins by virally encoded proteases is an effective viral strategy to modulate host processes and evade antiviral responses [3–5,45]. In enterovirus infection, targeting 14-3-3ε as well as several other factors in the RLR-signaling pathway are critical for productive infection and points to the importance of ensuring evasion of this pathway. It will be of interest to examine the roles of other 14-3-3 proteins that are cleaved under enterovirus infection. The recent advances in N-terminomics to identify host targets of viral proteases [3–5,45], both *in vitro* and in virus-infected cells, will provide insights into other antiviral factors that must be counteracted to promote virus infection.

## Materials and Methods

### Contact for reagent and resource sharing

Further information and requests for resources and reagents can be directed to Eric Jan.

### Cells and viruses

293T and A549 cells were cultured in Dulbecco’s Modified Eagle Medium (DMEM, Gibco 12100-046) supplemented with 10% fetal bovine serum and 1% penicillin/streptomycin at 37°C. A549 cells were kindly provided by Dr. Robert Hancock (University of British Columbia). Poliovirus (Mahoney type 1 strain; accession NC_002058.3) was generated from a pT7pGemPolio infectious clone. Poliovirus and coxsackievirus B3 stocks were propagated and titered in HeLa cells. Influenza A virus (H1N1) strain A/California/07/2009 was obtained from ATCC (VR-1894), propagated in MDCK cells, and titered with hemagglutination assay (Novus, Cat no. NBP3-05281). Sendai virus (Cantell strain) was obtained from Charles River (Mat.

10100774).

#### Plasmids and transfections

GFP and myc-tagged 14-3-3ε were generated as follows: a G-block containing the full-length 14-3-3ε protein (IDT) was cloned into pEGFP-C1 (Addgene #2487) using XhoI and BamHI restriction sites. To generate truncated mutants via site-directed mutagenesis, the following primers were used: 5’-CTCGAGCTATGGATGATCGAGAGGATCTG-3’ (for 5’ end amplification of all constructs), 5’-GGATCCTTACTGCATGTCTGAAGTCCATAG-3’ (3CN) and 5’-GCTCTTCTTAGTCACCCTGCATGTCTGAAGT-3’ (CaspN). To swap the GFP tag with a myc tag, a double-stranded oligo (5’-GCTAGCGCCGCCATGGTGGAGCAAAAGCTCATTTCTGAAGAGGACTTGAGATCT-3’) was subcloned into the pEGFP-C1 plasmid using restriction sites NheI and BglII. To generate scanning alanine mutants, DNA fragments containing the desired mutations were synthesized and subcloned into the parent vector (myc-WT 14-3-3ε) using XhoI and BamHI restriction sites.

All DNA transfections were performed using Lipofectamine 2000 according to manufacturer protocol (ThermoFisher, 11668019). Briefly, cells were seeded 24 hours before transfection to allow adherence. Transfection took place in antibiotic-free DMEM supplemented with 10% fetal bovine serum. All transfections were conducted for 16-24 hours prior to experimentation. For siRNA transfections, knockdowns were performed using DharmaFECT 1 (Dharmacon, T-2001-03) and a custom siRNA duplex (sense: 5’-CAUCUAAGAGAGAGGUUAAUU-3’; antisense: 5’-UUAACCUCUCUCUUAGAUGUU-3’) according to manufacturer protocol. When applicable, cells were transfected with 10 µg/mL polyinosinic:polycytidylic acid (poly(I:C); Invivogen).

### Virus infections

For poliovirus, coxsackievirus B3, and Sendai virus, virus was incubated with cells at the indicated MOI for 1 hour in DMEM + 1% penicillin/streptomycin at 37°C. After adsorption, media was replaced with complete DMEM (1X DMEM, 10% fetal bovine serum, 1% penicillin/streptomycin) and incubated for the designated time. For infections in the presence of zVAD-FMK (RD Systems, FMK001), zVAD-FMK was added to a final concentration of 50 µM 16 hours prior to infection and virus-containing media was supplemented with 50 µM zVAD-FMK for the duration of infection.

For Influenza A virus infections, cells were infected with H1N1 at the indicated MOI in viral growth medium (1X DMEM, 1% penicillin/streptomycin, 0.2% BSA, 25 mM HEPES pH 7.4, 0.5 µg/mL TPCK-treated trypsin) for 6 or 24 hours.

### Proteomic datasets

The TAILS dataset used for hypothesis generation was previously established and described in Jagdeo *et al* [3], and is publicly available in the ProteomeXchange Consortium (proteomecentral.proteomexchange.org) database under the accession number PXD008718.

### Immunoblot analysis

Equal amounts of protein were resolved on an SDS-PAGE gel and subsequently transferred onto a polyvinylidene difluoride (PVDF; Millipore) membrane. Primary antibodies used were as follows: 1:2000 α-Tubulin (ab4074); 1:1000 14-3-3ε (CST #9635); 1:1000 14-3-3 sampler kit (CST #9769T); 1:1000 RIG-I (CST #3743); 1:2000 c-Myc antibody (Thermo MA1-980); 1:2000 myc tag antibody (Bethyl A190-105A); 1:2000 FLAG M2 (Millipore F1804); 1:1000 VDAC1 (ab15895); 1:1000 TBK1/NAK (CST #3504); 1:1000 phospho-TBK1/NAK (CST #5483); 1:1000 hnRNP M (sc-134360); 1:3000 VP1 (Dako); 1:1000 H1N1 Nucleocapsid (ab104870); 1:1000 β-Actin (ab8224); 1:1000 PARP (CST #9542); 1:2000 GFP (Roche 11814460001).

### In vitro cleavage assay

*In vitro* cleavage assays were performed as described [4]. Briefly, lysates were resuspended in cleavage assay buffer (20 mM HEPES, 150 mM KoAC, 1 mM DTT) in the presence of protease inhibitors (ThermoFisher, cat. no. 78440). Lysates were incubated with wild type or catalytically inactive poliovirus 3C^pro^ (C147A) and the reaction was quenched after the indicated time using SDS-PAGE loading buffer.

### RT-qPCR

RT-qPCR was performed using total cellular cDNA. Briefly, cells were harvested in 1 mL Trizol (ThermoFisher, 15596018) and total cellular RNA was isolated according to manufacturer protocol. RT-qPCR was performed using NEB Luna^®^ Universal One-Step RT-qPCR Kit (#E3005L) using 20 ng total RNA. The following primers were used for analysis: GAPDH (5’-GGTGGTCTCCTCTGACTTCAACA-3’, 5’-GTTGCTGTAGCCAAATTCGTTGT-3’), IFNB (5’-TAGCACTGGCTGGAATGAGA-3’, 5’-TCCTTGGCCTTCAGGTAATG-3’), NA (H1N1 viral RNA; 5’-CCGCCATGGGTGTCTTTC-3’, 5’-TCCCTTTACTCCGTTTGCTCCATC-3’).

### Trypan Blue Exclusion

Following transfection, cells were collected and gently washed with sterile PBS prior to resuspension. Cells were then diluted 1:1 with Trypan Blue stain and the number of cells stained were counted using a Countess™ 3 Automated Cell Counter (ThermoFisher).

### Membrane fractionation

Cell lysates were separated into cytosolic or mitochondrial fractions using the BioVision Mitochondria/Cytosol Fractionation Kit (BioVision #K256) according to manufacturer protocol. Briefly, cells were lysed by passing cells through a 25g needle 25-30 times. Unbroken cells and nuclei were cleared by centrifugation at 700g until no pellet was observed. The samples were then spun at 10,000g and the supernatant was saved as the cytosolic fraction. The remaining pellet was resuspended in mitochondria extraction buffer as the mitochondrial fraction.

### Co-Immunoprecipitation

Cells were lysed in lysis buffer containing 1% Triton-X, 150 mM NaCl and 1X HALT protease inhibitors (ThermoFisher #78429) on ice for 20 minutes before clarification. 500 µg-1 mg of cell lysate was then incubated with 1:100 w/w myc tag antibody (Bethyl A190-105A) or anti-FLAG M2 (Millipore F1804) overnight at 4C. The next day, Pierce© Protein A/G Magnetic Beads (ThermoFisher, 88802) were pre-washed with lysis buffer and added to the lysate/antibody mixture for 2 hours at 4C. Beads were then separated using a magnetic stand and washed twice with ice-cold lysis buffer (NaCl adjusted to 300 mM) and twice with ice-cold dH_2_O. Antibody-bound protein was then eluted from the agarose beads using sample buffer at room temperature for 10 minutes.

### Plaque Assay

Plaque assays were performed as follows: a confluent monolayer of 293T cells were infected with a dilution series (10^−1^ to 10^−7^) of virus absorbed in a minimum volume of serum-free media for 1 hour. Inoculum was subsequently removed and the monolayer overlaid with 1% methylcellulose in DMEM for 72 hours. The overlay was then discarded and the cells were stained for 15 minutes at room temperature in 1% crystal violet (w/v) and 50% methanol (v/v).

### Statistical analysis

All graphs and statistical analyses were created using GraphPad Prism 9.0. **\* =** p>0.05; ** = p>0.005; *** = p>0.0005; **** = p>0.00005. For RT-qPCR data, error bars represent +/-95% confidence interval; for all other statistical analyses, error bars represent the standard deviation of at least n=3 biological replicates.

## Acknowledgements

The authors thank members of the Jan lab for helpful input and discussion. The authors thank Dr. Robert Hancock (UBC) for A549 cells and Dr. Sean Workman (University of Regina) for computational modeling and help with statistical analysis. This study was supported by the Canadian Institutes of Health Research (E.J: PJT-156351, H.L: PJT-173318, C.M.O: FDN-148408) and a NSERC Discovery Grant (E.J: RGPIN-2017-04515, H.L: RGPIN-2022-02979). D.D.T.A. was supported by a National Sciences and Engineering Research Council (NSERC) Doctoral Postgraduate Scholarship and a UBC Four-Year Fellowship.

## Author Contributions

Conceptualization: D.D.T.A, Y.M, H.L, C.M.O, E.J; Investigation: D.D.T.A, Y.M, M.V, D.A.B, B.N.H, E.J.; Formal Analysis: D.D.T.A, E.J.; Methodology: D.D.T.A, Y.M, H.L, C.M.O, E.J; Validation: D.D.T.A, Y.M; Supervision: L.J.F, H.L., C.M.O, E.J.; Writing-original draft: D.D.T.A, E.J; Writing-review and editing: D.D.T.A, Y.M, M.V, D.A.B, B.N.H, L.J.F, H.L, C.M.O, E.J; Funding Acquisition: L.J.F, H.L, C.M.O, E.J.

